# Unified access to up-to-date residue-level annotations from UniProt and other biological databases for PDB data via PDBx/mmCIF files

**DOI:** 10.1101/2022.08.10.503473

**Authors:** Preeti Choudhary, Stephen Anyango, John Berrisford, Mihaly Varadi, James Tolchard, Sameer Velankar

## Abstract

More than 58,000 proteins have up-to-date correspondence between their amino acid sequence (UniProtKB) and their 3D structures (PDB), enabled by the Structure Integration with Function, Taxonomy and Sequences (SIFTS) resource. In addition to this fundamental mapping, SIFTS incorporates residue-level annotations from other biological resources such as Pfam, InterPro, SCOP, SCOP2, CATH, IntEnz, GO, PubMed, Ensembl, NCBI taxonomy database and Homologene. The SIFTS data is exported in XML format per individual PDB entry and is also accessible via the PDBe REST API. These mappings have always been maintained separately from the structure data (PDBx/mmCIF file) in the PDB archive. In this current work, taking advantage of the extensibility of the core PDBx/mmCIF framework, we extended the wwPDB PDBx/mmCIF data dictionary with additional categories to accommodate SIFTS data and added the UniProt, Pfam, SCOP2, and CATH mapping information directly into the PDBx/mmCIF files from the PDB archive. The integration of mapping data in the PDBx/mmCIF files provides consistent numbering of residues in different PDB entries allowing easy comparison of structure models. The extended PDBx/mmCIF format yields a more consistent, standardised metadata description without altering the core PDB information. This development enables up-to-date cross-reference information at residue level resulting in better data interoperability, supporting improved data analysis and visualisation.

**Availability and implementation:** We expanded the PDBe release pipeline with a process that adds SIFTS annotations to the PDBx/mmCIF files for individual structures in the PDB archive. The scientific community can download these updated PDBx/mmCIF files from the PDBe entry pages

(https://pdbe.org/7dr0) and through direct URLs

(https://www.ebi.ac.uk/pdbe/static/entry/7o9f_updated.cif), using the PDBe download service

(https://www.ebi.ac.uk/pdbe/download/api) or from the EMBL-EBI FTP area

(https://ftp.ebi.ac.uk/pub/databases/msd/updated_mmcif/).

## Introduction

As of July 2022, the Protein Data Bank (PDB)^1^ contains over 190,000 entries representing over 58,000 unique entries in the Universal Protein Resource (UniProt)^2^. Often, the PDB archive has the same protein in multiple entries under different experimental conditions or interacting with different macromolecules (proteins, DNA, RNA) or ligand molecules^3–5^. Multiple 3-dimensional coordinates of the same protein are invaluable for comparative structure-function studies^3,6,7^. Linking structure data with annotations available in other data resources such as UniProt^2^ and to the structural and functional annotations is critical in order to understand biological function and processes at molecular level. However, one of the barriers to comparative analysis or data integration is the independent, depositor-provided residue numbering in the coordinate files, which may not be the same as the protein sequence numbering^8^. While solving a protein 3D structure, many times the experiments are carried out only on a part of complete protein molecules (e.g. a domain) to either make the sample amenable to experimental methods especially in cases where there are highly flexible linker regions or intrinsically disordered regions^9,10^. Around 58% of the structures in the PDB contain smaller fragments (e.g. a domain) corresponding to different regions of a protein sequence. To determine where these fragments are located on the full-length protein sequence, these fragments need to be mapped to a common reference e.g. protein sequence numbering in UniProt database. The situation becomes complicated as often the flexible regions in the protein molecules are not modelled and hence there are unobserved residues in protein structures. The occurrence of missing residues makes structure-to-sequence mapping even more challenging.To address this fundamental problem of standardising residue numbering to make protein structure data more accessible to the broader scientific community, the PDBe^11^ and UniProtKB^2^ teams collaborated to establish the Structure Integration with Function, Taxonomy and Sequences (SIFTS) resource in 2002^12,13^. SIFTS provides up-to-date residue level mapping between UniProt protein sequences and PDB protein structures allowing better integration of annotations based on protein sequence and structure.

In addition to mapping PDB structures to UniProt sequences, SIFTS also maps to other biological resources such as Pfam^14^, InterPro^15^, SCOP^16^, CATH^17^, IntEnz^18^, GO^19,20^, Ensembl^21^, NCBI taxonomy database^22^ and Homologene^23^.

In the past 20 years, SIFTS has become an essential resource, and its data provides the foundation of many data services and web pages. SIFTS is fundamental to the PDBe and PDBe-KB data resources^24^ and other databases, such as UniProt^2^, Pfam^14^, RCSB PDB^25^, PDBj^26^, SCOP2^27^, InterPro^15^ and MobiDB^28^, rely on SIFTS to fetch cross-references between PDB structures and other biological databases. SIFTS data is distributed as summary flat files in CSV/TSV formats and also as a detailed per-entry XML files with residue-level information available from the EMBL-EBI FTP area (ftp://ftp.ebi.ac.uk/pub/databases/msd/sifts/). SIFTS data is also accessible via the PDBe API^29^.

While SIFTS data has significantly improved interoperability of PDB structure data with other key data resources, it still requires to be accessed separately from the 3D coordinates data in the PDB. The SIFTS output format is incompatible with 3D visualisation software and requires an additional step of parsing the data to display SIFTS annotations on protein 3D structure. To make further progress in the usability and interoperability of PDB structures, the next logical step is to integrate SIFTS annotations next to the 3D coordinates in the PDBx/mmCIF files. Moreover, with the availability of numerous high-quality protein structure models from resources like SWISS-MODEL^30^ and AlphaFold DB^31,32^, which generally follow the protein sequence numbering scheme, it was timely and essential to augment the protein sequence numbering for the experimentally determined 3D coordinates in the PDB. Using data from SIFTS resource, PDBrenum^8^ web server replaces author sequence numbering with UniProt numbering in PDB or PDBx/mmCIF format files but it has certain limitations while handling special cases. For instance, while renumbering if this web server does not find any mapping in SIFTS, it simply adds a large number to the residue’s sequence position number. These residues can be expression tags or insertions and need to be represented appropriately without losing experimental context of the sample. Similarly, for chimeric proteins which are mapped to more than one protein sequence (UniProt accession), PDBrenum only renumbers according to the one protein sequence which has maximum coverage, losing information about remaining proteins in the chimeric construct. It does not integrate annotations to other data resources from SIFTS like Pfam, SCOP2 and CATH as well. Thus, there is a need to find a more consistent, sustainable and up-to-date solution while incorporating UniProt numbering and annotations from various other data resources in the 3D coordinate files.

Here, we describe our approach for incorporating SIFTS annotations in extended PDBx/mmCIF files so that the UniProt residue numbering is directly embedded next to the atomic coordinates. The PDBx/mmCIF is an extensible format that also provides mechanism to maintain data integrity and is the master format for macromolecular structure data in the PDB^33^. We describe how this current work extends the PDBx/mmCIF dictionary by leveraging the extensibility of its structured framework, thereby providing a mechanism to enrich the biological context of a PDB structure.

## Results

### Updated SIFTS process

SIFTS is a two-fold process^13^ which includes 1) a semi-automated process to retrieve the manually curated UniProtKB cross-reference (or canonical UniProtKB accession) for each protein chain in the PDB and 2) an automated process that generates residue-level correspondences between structure (PDB) and the corresponding sequence (UniProt). Initial mapping of UniProtKB sequence to the PDB structure is manually curated during the wwPDB annotation process^34^. During the semi-automated process, these manually curated mappings are checked for obsoleted or secondary UniProt accessions and are updated accordingly. In the automatic process, the manually curated canonical accession is then expanded to include all its isoforms, and sequence alignment is computed for each PDB-UniProt pair. Taking only the PDB-UniProt pairs with the same source organism or atleast having a common ancestor within one or two levels up to species level in the taxonomy tree and having at least 90 % sequence identity, the pair with highest sequence identity is annotated as the best mapping. Once we have established the mapping between UniProt and PDB protein residues, the cross-references from other resources such as Pfam^14^, InterPro^15^, SCOP^16^, CATH^17^, IntEnz^18^, GO^19,20^, Ensembl^21^ and Homologene^23^ are added. The SIFTS annotations are stored in the SIFTS database, which is used to expose the data via the PDBe REST API. Individual XML files for each PDB entry with residue-level information are exported and the summary files are generated in CSV/TSV formats. An additional process reads the data from the SIFTS database and augments the PDB structure files with UniProt numbering and structure (SCOP2, and CATH resource) and sequence (Pfam resource) domains annotations. We refer to these extended files as updated PDBx/mmCIF files. This update yields more consistent, standardised metadata. It is important to note that none of the core PDB information, such as atomic coordinates and experimental data, are altered in any way. Figure 1 shows the schematic overview of the data flow of the updated SIFTS process. The process helps researchers and data services access SIFTS data directly from the updated PDBx/mmCIF^35^ files. To facilitate this update, additional “SIFTS-specific” mmCIF data categories were designed that extend the core PDBx/mmCIF data dictionary. These format specifications are discussed in detail below.

**Figure 1.**
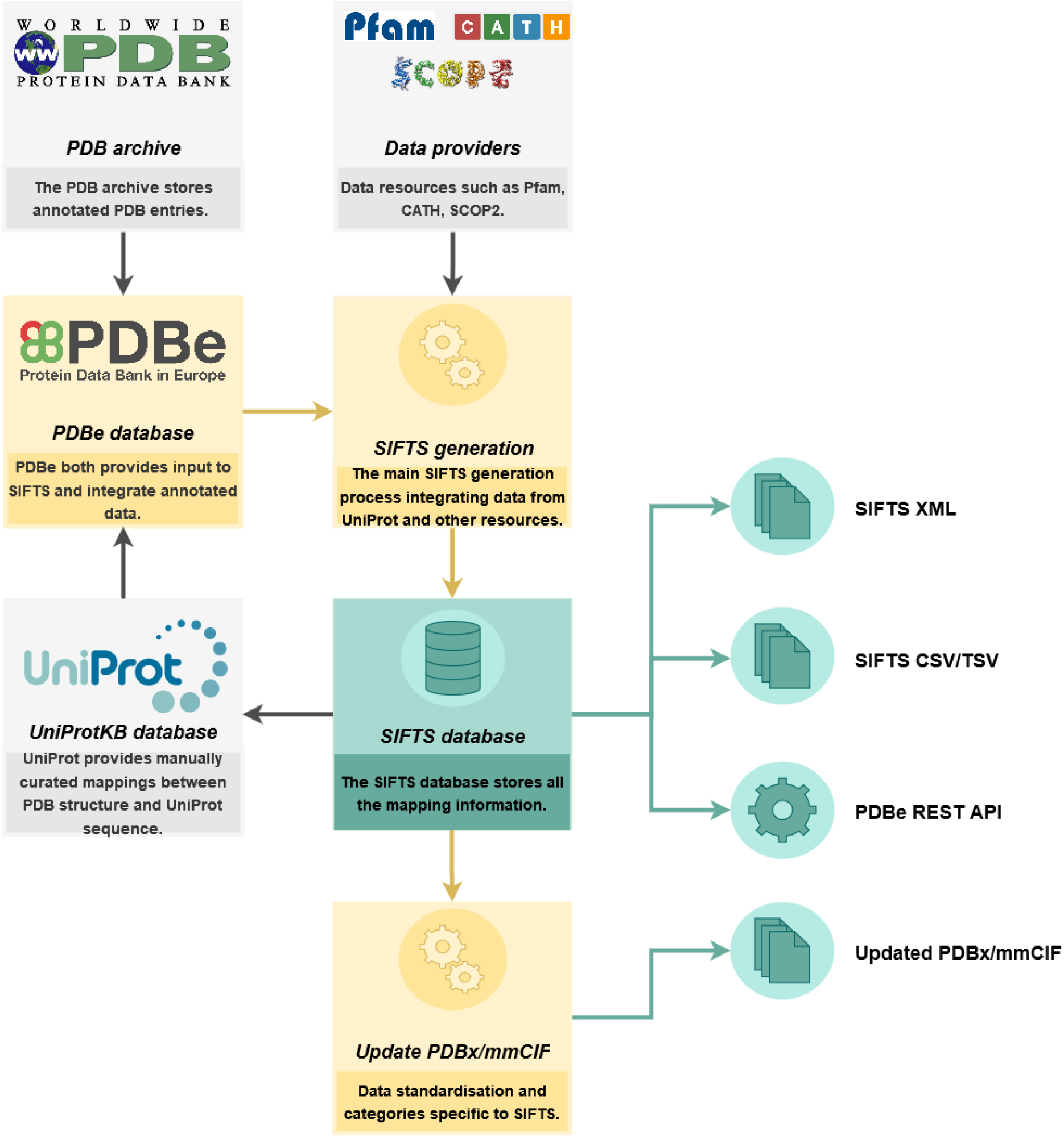
The overall data flow of the updated SIFTS process. The figure shows the components of the SIFTS process in yellow and the output of the SIFTS process in green. The process generates the SIFTS database and XML, CSV, TSV, and updated PDBx/mmCIF files. Components coloured in grey are data resources external to SIFTS.

### Format specification changes

PDBx/mmCIF framework organises information in categories containing related data items^35^. The updated PDBx/mmCIF files contain the residue mappings between UniProt, PDB, and annotations from Pfam, SCOP2, and CATH. There are two types of mappings: for a range of residues (per-segment) and for individual residues (per-residue). New data categories were added to represent these additional per-segment and per-residue mappings (Figure 2). Two new categories “_pdbx_sifts_unp_segments” and “_pdbx_sifts_xref_db_segments” were added to represent per-segment mapping to UniProt-KB and other data resources - Pfam, SCOP2, CATH respectively. A third category, “_pdbx_sifts_xref_db”, was added to provide residue-wise mapping from all the external resources. The “_atom_site” category, which represents the coordinate information, was extended with additional data items to integrate UniProt residue numbering from the best mapping next to the atomic coordinates.

**Figure 2.**
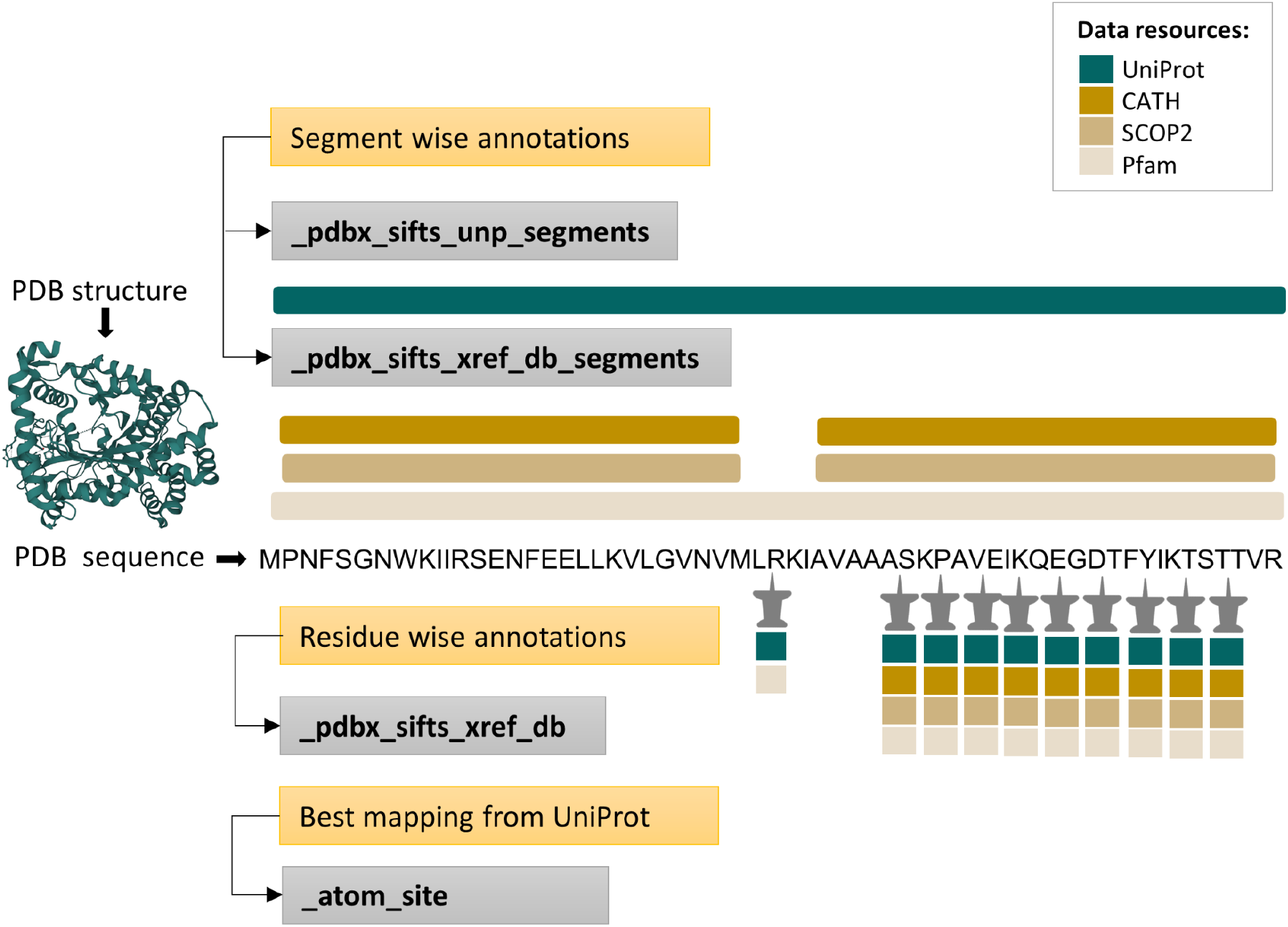
The PDBx/mmCIF extension incorporates mappings from various data resources. SIFTS annotations mapping PDB residues to various data resources are shown both segment-wise (top) and residue-wise (bottom). All the new SIFTS-specific or modified PDBx/mmCIF categories are shown in grey boxes. The new SIFTS-specific PDBx/mmCIF categories introduced to show segment-wise annotations from UniProt-KB and all the other external data resources(Pfam, SCOP2, CATH) are “_pdbx_sifts_unp_segments” and “_pdbx_sifts_xref_db_segments” respectively. “_pdbx_sifts_xref_db” is another new SIFTS-specific PDBx/mmCIF category introduced to show residue-wise annotations. We also modified the “_atom_site” category to indicate the best mapped UniProt sequence.

A summary of the new and modified data categories necessary to encode the SIFTS annotations data is provided below :

1. **_pdbx_sifts_unp_segments** This new category describes residue range-based cross-references specific to the UniProt database. It shows segments/regions of PDB residues mapped to the canonical UniProt accession and all its isoforms. The residue mapping is established by aligning the PDB sequence to each UniProt accession (canonical and all the isoforms) and the sequence identity between the aligned PDB-UniProt pair is provided. This category also indicates the best mapped UniProt accession.
2. **_pdbx_sifts_xref_db_segments** This new category describes residue range-based cross-references to additional databases such as Pfam, SCOP2, and CATH.
3. **_pdbx_sifts_xref_db** PDB structures often have missing residues, expression tags or linker regions, making the expansion of mappings from segments (residue-range) to individual residues cumbersome. An essential category, “_pdbx_sifts_xref_db”, therefore describes residue level cross-references to external databases. This category provides annotations specific to the best mapped UniProt accession and can be used to identify all the mappings for each individual residue to external databases (Figure 3).
4. **_atom_site** New data items were added to the “_atom_site” category to represent the best mapped UniProt accession, residue type and number. The new data item “_atom_site.pdbx_label_index” provides a unique identifier for all the polymer residues and individual non-polymer and solvent components.

**Figure 3.**
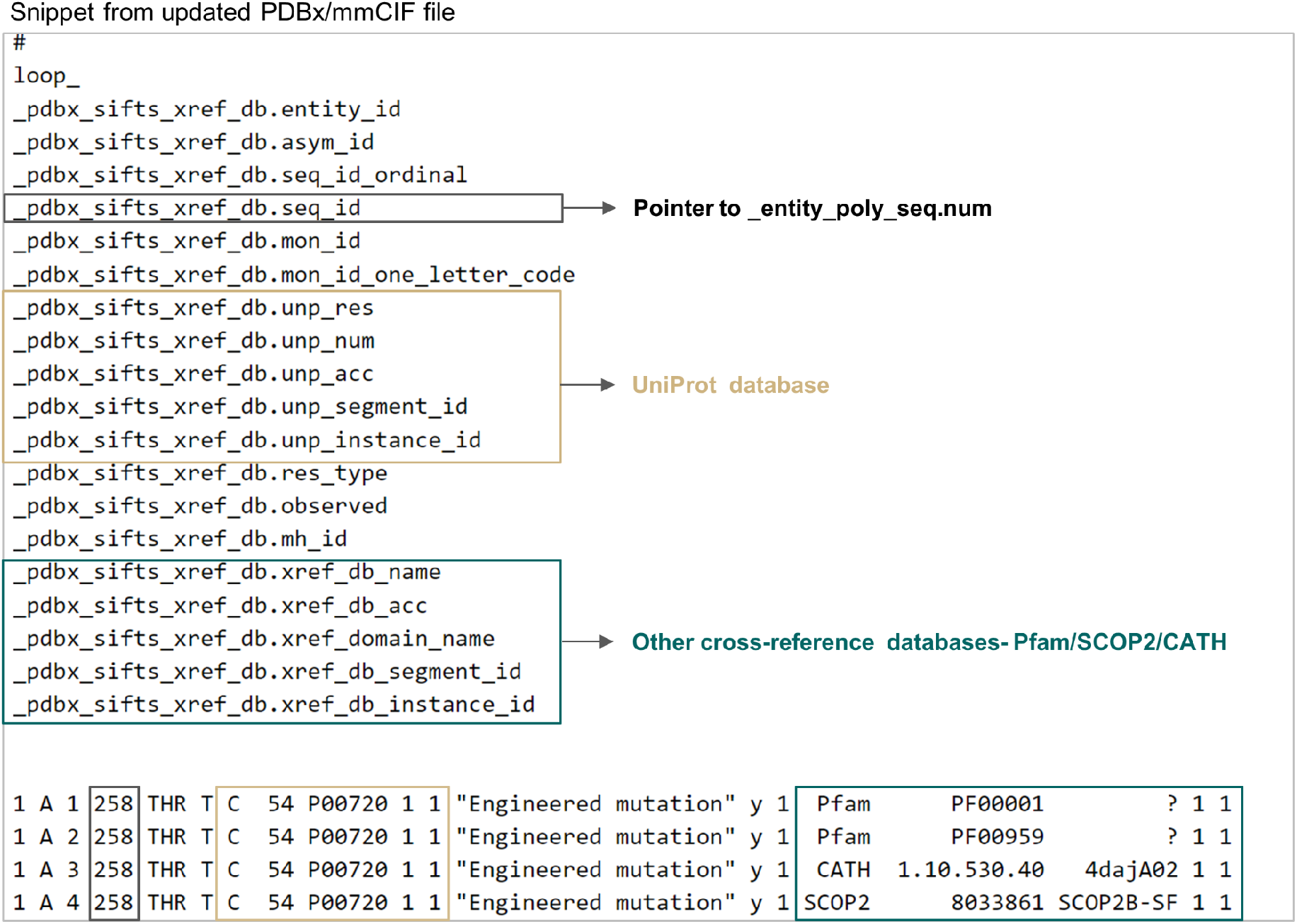
Single placeholder in PDBx/mmCIF files to find all the annotations associated with any residue from external databases. This figure shows the “_pdbx_sifts_xref_db” category for PDB entry 4daj. This critical new data category can describe residue-level cross-references to external databases. The items specific to the UniProt database and other cross-reference databases are marked in beige and green colored boxes respectively.

There are two different numbering schemes followed to indicate each residue (amino-acid or nucleotide) in the PDBx/mmCIF file. Firstly, “auth_seq_id” which is the numbering provided by the author. Author can assign its value in any desired way and the values may be used to relate the given structure to a numbering scheme in a homologous structure, including sequence gaps or insertion codes, which are not necessarily numbers. Secondly, “label_seq_id” which is the wwPDB assigned numbering which starts from 1 and increments sequentially for all the polymer residues. All the SIFTS-specific categories refer consistently to the wwPDB assigned numbering scheme defined by the “label_seq_id” data item in the atom_site category. The reference to labl_seq_id is provided by the data items “.seq_id”, “.seq_id_start” and “.seq_id_end” in the relevant categories. Data on the author provided or the PDB numbering scheme can be retrieved using the appropriate relationships defined in the PDBx/mMCIF categories(Figure 4).

**Figure 4.**
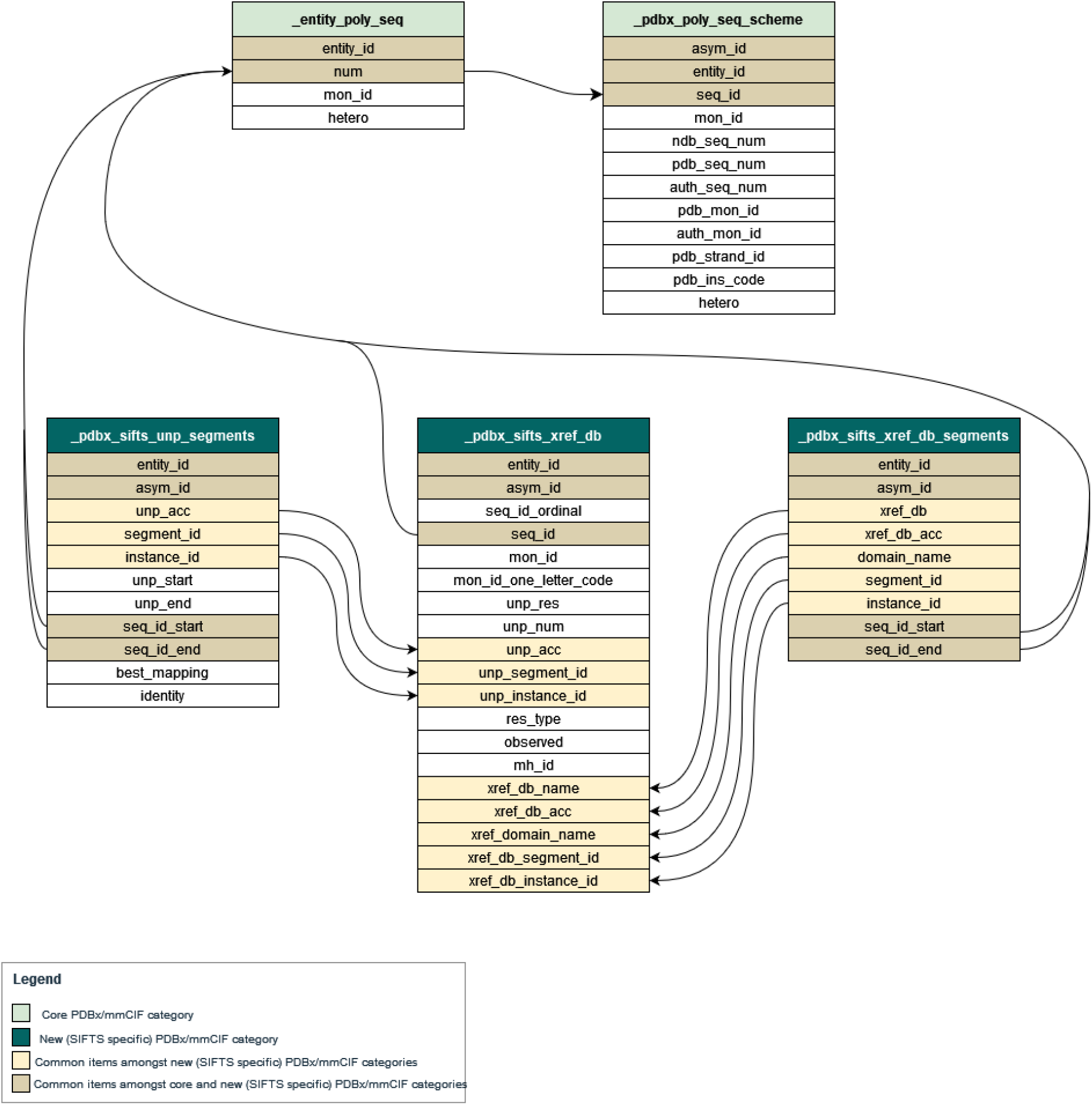
Category relationship diagram including new SIFTS specific PDBx/mmCIF categories. New SIFTS specific PDBx/mmCIF data categories are shown along with their data items. All the common data items amongst these new data categories are highlighted and their relationship is shown. Further, the relationship of data items representing PDB residue numbers - “.seq_id”, “.seq_id_start” or “.seq_id_end” in these new data categories to core data categories is shown.

Often in many proteins several domains are tandemly repeated^36^. Additionally, researchers also synthesise structures where even the entire protein is repeated for specific research purposes^37,38^. Previously, there was no automated way to find corresponding UniProt mappings for various copies of such protein structures in the PDB. We designed the data item “.instance_id” to help identify multiple instances of the same protein segment. For example, in the single-chain dimeric Streptavidin structure (PDB 6s50), the two copies of Streptavidin^39^ are easily identified by instance ids “1” and “2” for the UniProt accession P22629 (Figure 5).

**Figure 5.**
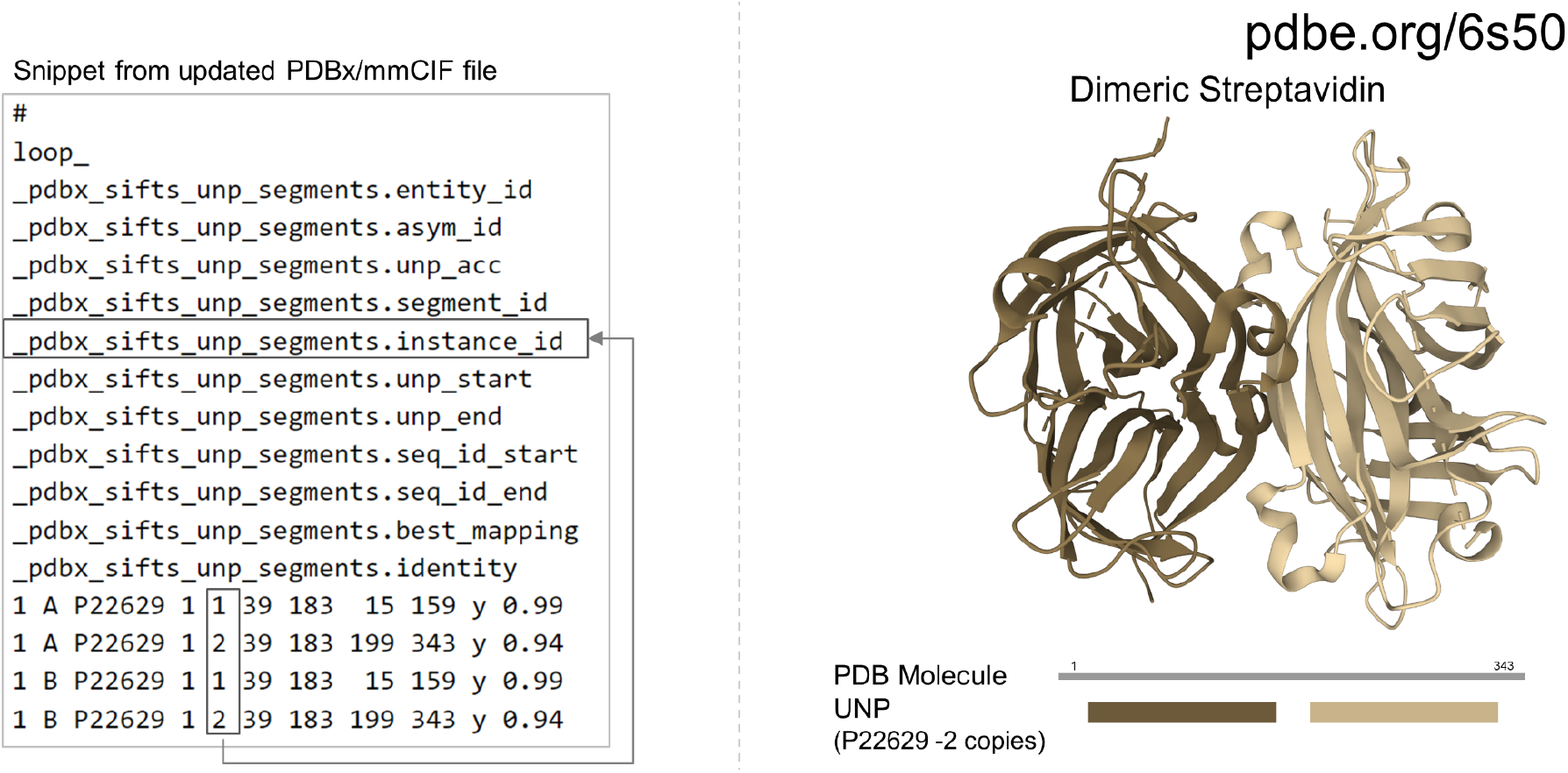
Distinguishing between multiple instances of the same protein in the updated PDBx/mmCIF file. The data item “.instance_id” enables scientific users to identify the two copies of the same protein, Streptavidin (UniProt accession P22629), in the dimeric Streptavidin structure (entry 6s50).

Similarly, users can rely on this data item to easily identify multiple copies of the same domains in a protein structure.

During evolution protein structures may evolve with an insertion of an additional domain which splits the original structural domain into a discontinuous range of residues in the sequence^40^. For example, the *E*.*coli* enzyme RNA 3’-terminal phosphate cyclase (PDB 1qmh) consists of two domains where a smaller insert domain (residues 186-276) splits the larger (parent) domain^41^. The identification of the split parent domain (residues 5–182, 277–337) is evident from the “.segment_id” data item (Figure 6).

**Figure 6.**
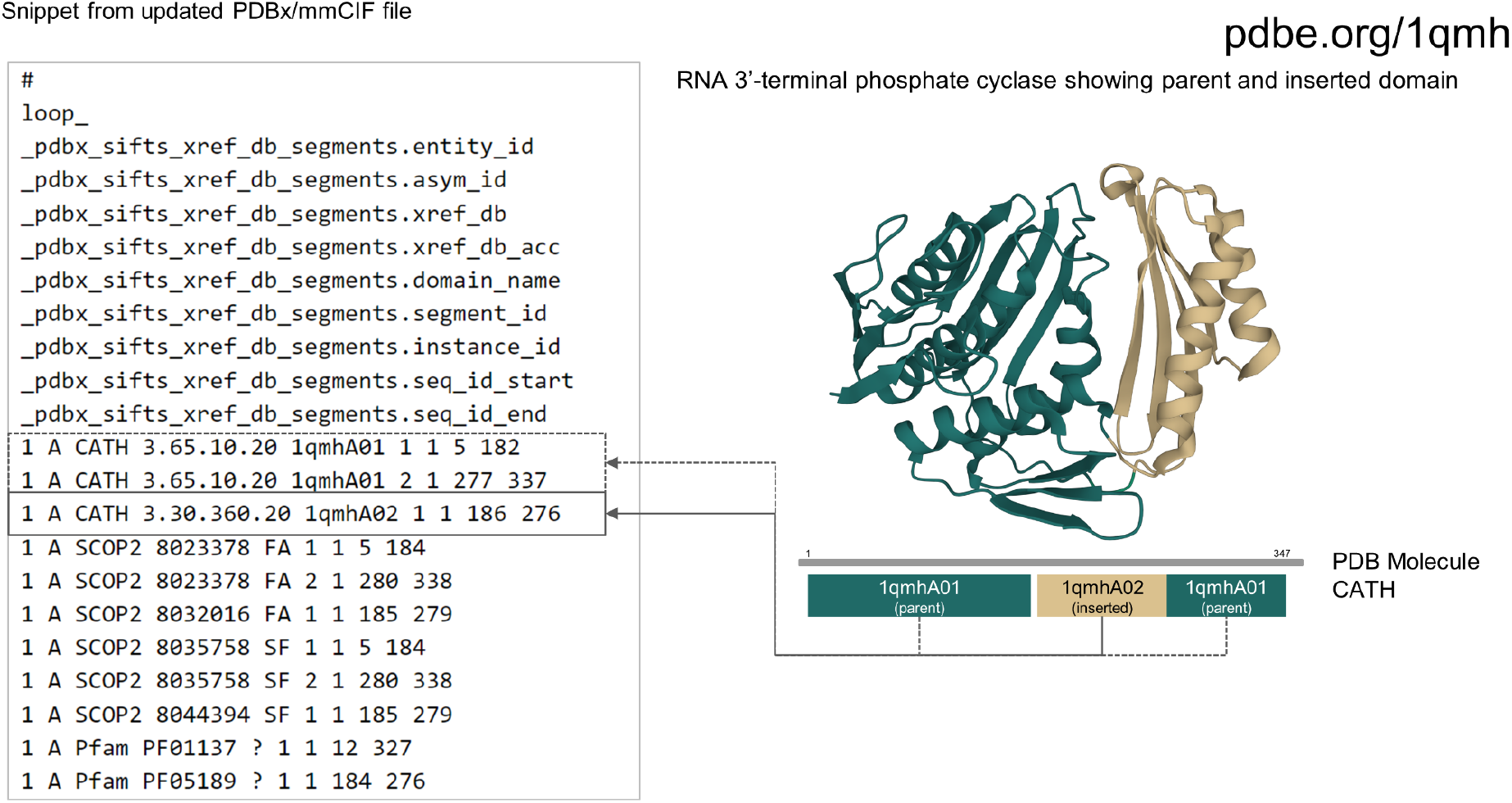
Identification of split domains from PDBx/mmCIF file. The “_pdbx_sifts_xref_db_segments” category in the PDBx/mmCIF file of PDB 4daj helps to clearly identify discontinuous domains. The two halves of the M3 receptor domain are indicated by the same “.instance_id” but different “.segment_id”.

Complete documentation for all the new and updated data categories and items is available at https://mmcif.wwpdb.org/dictionaries/mmcif_pdbx_v5_next.dic/Index/.

## Application

SIFTS data has been used in many research studies to retrieve residue correspondence between PDB structures and UniProt sequences^42–46^. Often authors have to manually renumber the coordinate file to reflect UniProt numbering for subsequent comparative analysis^47–49^. SIFTS resource has also been used to fetch annotations like sequence domains and structural domains for various PDB structures^49,50^. The updated PDBx/mmCIF files with SIFTS annotations introduced in this work addresses this fundamental need to combine data from various resources and provide coordinate files with a common reference frame helping researchers to perform such comparitive analysis more efficiently.

Various data visualisation tools can directly use these updated PDBx/mmCIF files, making the mapping of 1D data onto the 3D views simpler. With our improvements, researchers from various scientific fields can easily map sequence feature data onto PDB structures. For example, using these updated PDBx/mmCIF files, variant data can easily be mapped onto PDB structures. Users can directly retrieve all the SIFTS annotations like structural domains, sequence domains and conflicts between sequences and structures from these updated PDBx/mmCIF files.

These files also provide a basis for improved comparisons between experimentally determined and theoretical predicted protein models. UniProt numbering in the coordinate files allows direct residue correspondence making structural comparison and superposition easier. It also makes it easier to compare PDB structures with the predicted model structures from AlphaFold DB^31,32^, SWISS-MODEL^30^, RoseTTAFold^51^, and many other resources, as these models follow a natural sequence numbering. These files are already being used by Mol*^52^ (https://molstar.org/viewer/) to perform extremely fast superpositions using the SIFTS UniProt mapping. This superposition functionality in Mol*^52^ is very powerful as it gives users the means to directly superimpose protein structures in their web browser without downloading any data or software. Figure 7 shows the superposition of the unbound and bound forms of human PTP1B protein (Protein Tyrosine Phosphatase 1B, UniProt accession: P18031) performed using the UniProt button (highlighted in red box) in the Superposition panel in Mol*^52^. This protein is known to be a signalling molecule regulating a variety of cellular processes including cell growth, differentiation and oncogenic transformation and is a potential therapeutic target for treatment of type 2 diabetes and cancer^53^. Upon substrate/inhibitor binding, the WPD loop transitions from an open to a closed conformation^54–57^ as shown in figure 7.

**Figure 7.**
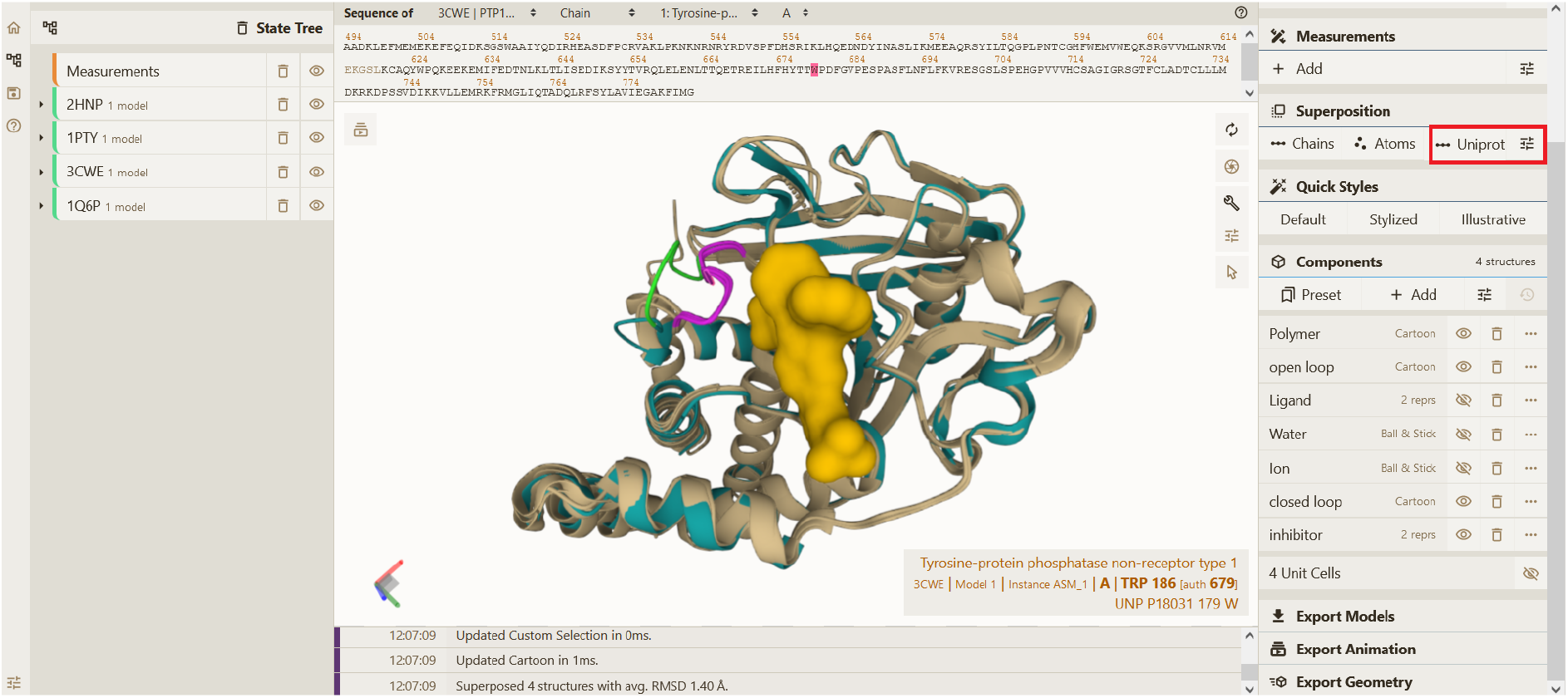
Superposition of protein structures using Mol*. The superposed apo and holo forms of human PTP1B protein are shown in green and beige colour, respectively, in Mol*. The WDP loop is in open (light green colour) conformation in the apo form (PDB ID 2HNP). Upon binding to various substrates/inhibitors this WDP loop attains closed (pink colour) conformation covering the catalytic site. The inhibitor bound in PDB ID 1Q6P is shown in surface representation. The average RMSD between the four superposed structures as computed by Mol* is 1.40 Å. As seen in tool-tip (bottom-right in figure), residue W179 from PDB ID 3CWE and other residues in inhibitor bound PDB entries-3CWE and 1Q6P have different author numbering compared to the unbound/substrate bound form (PDB ID 2HNP/1PTY). The UniProt numbering in the updated PDBx/mmCIF file provides a common reference frame for residue correspondence and supports superposition based on Uniprot in Mol*.

The new, updated PDBx/mmCIF files also provided a basis for developing interactive visualisations. For example, on the PDBe entry pages, we now show the ProtVista component, a 2D visualisation for displaying the primary sequence features of proteins. ProtVista was developed in collaboration with UniProt and InterPro at EMBL-EBI. With the help of the updated PDBx/mmCIF files, ProtVista (Figure 8A) could easily be linked to a 2D topology component (Figure 8B) and the Mol* 3D viewer (Figure 8C). As shown in Figure 8, for Mannose-1-phosphate guanyltransferase, PDB entry 7d72 (https://www.ebi.ac.uk/pdbe/entry/pdb/7d72/protein/1), if users click on any residue annotation in the 2D viewer ProtVista, it is automatically highlighted that residue in 3D, in the Mol* viewer. Similarly, users can highlight various structural or sequence domains, or other annotations in either the 2D topology component, 2D ProtVista component or Mol* viewer, and the three visualisations cross-talk with each other simultaneously, making visualisation and interpretation of data much easier. Mol* already uses these updated PDBx/mmCIF files to display various annotations on PDBe and PDBe-KB webpages. With SIFTS annotations directly available in the coordinate file, the 3D visualisation on PDBe and PDBe-KB webpages is more efficient and optimal.

**Figure 8.**
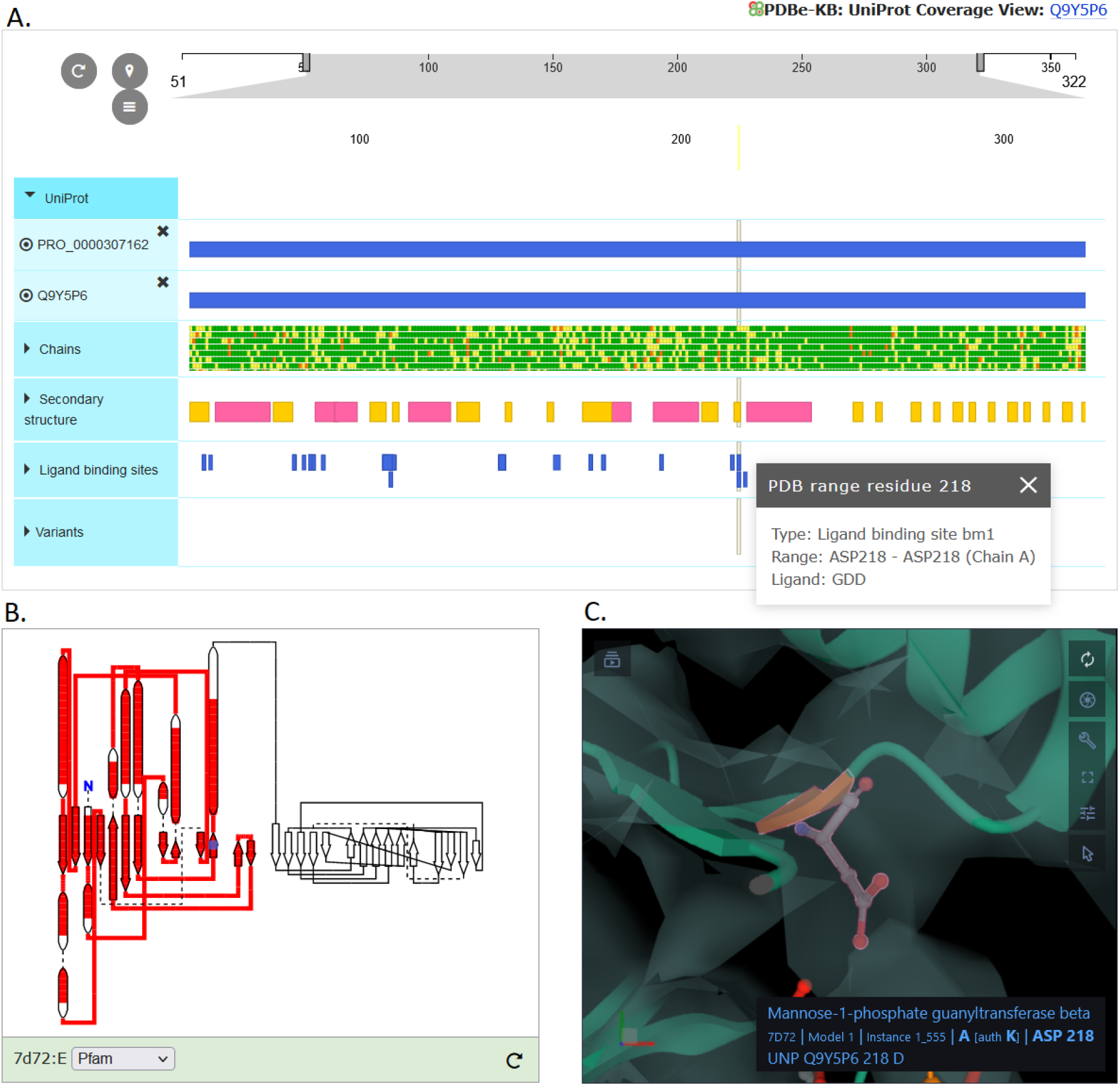
The 2D visualisation components are interactively linked with 3D visualisation components on PDBe entry pages. Various 2D and 3D visualisation components seen on PDBe entry pages are interactively linked with each other. Here we show visualisation data for Mannose-1-phosphate guanyltransferase (PDB ID 7d72). A. shows 2D sequence feature viewer (ProtVista) and B. shows 2D topology viewer, along with C. showing the 3D viewer, Mol*. As users select any residue (here ligand-binding residue ASP218 is selected) in ProtVista, it is automatically highlighted in Mol* and vice-versa. Users can also highlight Pfam domains in any of these viewers. Here, we show the Pfam domain highlighted in red in the 2D topology viewer.

## Conclusion

Interoperability challenges between the protein structure data in PDB and the protein sequences in UniProt presents a significant barrier to accessibility and usability. The seemingly trivial task of mapping residue-level information proved to be a formidable task that necessitated the development of the SIFTS resource. While SIFTS has successfully provided up-to-date mappings between the PDB and other data resources for the past 20 years, using these mappings still required some level of technical expertise.

To remove a tedious but previously mandatory step in many structural data analyses, we worked on adding the SIFTS mapping data directly into the PDBx/mmCIF files, the master format for the PDB archive. We designed new data categories and extended existing ones to provide flexible support for residue-level annotations. This development will allow easy linking of structural and functional annotations derived using structure and sequence data. It will also streamline the vast majority of high-throughput bioinformatics analysis pipelines by allowing developers to remove a tedious and error-prone step from their processes. Including the SIFTS data in the updated PDBx/mmCIF will also improve the efficiency of data visualisation tools, both those that specialise in 3D molecular graphics and those that focus on the interactive mapping of annotations onto to the protein structure representations e.g. sequence or topology.

By extending the PDBx/mmCIF data format, this work has laid the foundation for the future integration of additional annotations, allowing the files to be more comprehensive and to provide the biological context for PDB structures.

## Acknowledgement

This paper is dedicated to the fond memory of our dear collaborator and wwPDB member John Westbrook. John critically reviewed the new “SIFTS-specific” categories in PDBx/mmCIF data dictionary and provided valuable feedback. We also thank Ezra Peisach for his valuable comments while updating the PDBx/mmCIF data dictionary. Both John Westbrook and Ezra Peisach are members of the RCSB Protein Data Bank, a co-founder of the wwPDB along with PDBj and PDBe. We also thank EMBL and EMBL-EBI for providing the necessary infrastructure to run this process weekly.

## Funding

This work was supported by funding from EMBL and funding awarded to PDBe by the UK Biotechnology and Biological Research Council (BB/V004247/1, PI:Sameer Velankar) and RCSB PDB by the NSF (DBI-2019297, PI: S.K. Burley) supporting development of a Next Generation PDB archive.

## Conflict of interest

None declared

## Notes

### Competing Interest Statement

The authors have declared no competing interest.

